# Bio-hybrid Soft Robotic Bioreactors for Mimicking Multi-Axial Femoropopliteal Artery Mechanobiology

**DOI:** 10.1101/2021.09.24.461639

**Authors:** Cody Fell, Trent L Brooks-Richards, Maria Ann Woodruff, Mark C Allenby

## Abstract

The emerging field of soft robotics aims to emulate dynamic physiological locomotion. Soft robotics’ mimicry of naturally complex biomechanics makes them ideal platforms for exerting mechanical stimuli for patient-specific tissue maturation and disease modeling applications. Such platforms are essential for emulating highly flexible tissues such as the kneecap’s femoropopliteal artery (FPA), one of the most flexible arteries in the body, which flexes and bends during walking, standing, and crouching movements. The FPA is a frequent site of disease, where 80% of all peripheral artery diseases manifest, affecting over 200 million people worldwide. The complex biomechanical and hemodynamic forces within the FPA have been implicated in the frequent occurrence of PAD and lead to debilitating morbidities, such as limb-threatening ischemia. To better mimic these complex biomechanics, we developed an *in-vitro* bio-hybrid soft robot (BSR). First, Platsil OO-20 was identified as an ideal hyperelastomer for both cell culture and BSR fabrication using 3D printed molds. Then, employing a simulation-based design workflow, we integrated pneumatic network (PneuNet) actuators cast with Platsil OO-20, which extend in angular, longitudinal, and radial dimensions. Pressurizing the BSR PneuNets enabled a range of mechanical stimuli to be dynamically applied during tissue culture to mimic normal and diseased FPA flexions during daily walking and sitting poses, the most extreme being radial distensions of 20% and angular flexions of 140°. Finally, these designed, manufactured, and programmed vascular BSRs were seeded with mesenchymal stem cells and conditioned for 24 hours to highlight the effect of dynamic conditioning on cultured cell alignment, as well as type IV collagen production and the upregulation of smooth muscle phenotypes. Soft robotic bioreactor platforms that accurately mimic patient-, disease-, and lifestyle-specific mechanobiology will develop fundamental disease understanding, preoperative laboratory simulations for existing therapeutics, and biomanufacturing platforms for tissue-engineered implants.

## 1. Introduction

Soft robotics is a relatively new field of bioinspired robotics that aims to emulate physiologic locomotion. The movement of pneumatic network (PneuNet) actuated soft robots is controlled by pressurizing intelligently designed networks incorporated within an elastomer body to induce gradual actuation [1]. Recent examples of bioinspired PneuNets mimic the complex biomechanics of anatomical systems, such as fingers or octopus tentacles [2, 3]. In addition, devices designed to interface with biology and medicine are underway, from biohybrid robots that actuate via contractile cell tension to implantable soft robots designed to support heart function [4-6]. Characteristically composed of soft elastomeric materials that are often biocompatible, soft robots are an enabling technology for advanced bioreactors that mimic the complexities of soft tissue mechanobiology.

Bioreactor platforms that impart mechanical stimuli on growing cells *in vitro* enable more functional tissue regeneration when compared to static cultures. For example, tissue-engineered vascular grafts (TEVGs) matured with cyclic radial stretch have improved cell organization and higher extracellular matrix (ECM) deposition and alignment [7, 8]. Furthermore, cells respond differently when exposed to different modes, frequencies, and magnitudes of mechanobiological stimuli. For example, mesenchymal stem cells (MSCs) exposed to shear stress express endothelial phenotypes, whereas MSCs exposed to uniaxial stretch but not equiaxial stretch express vascular smooth muscle cell (SMC) phenotypes [9-11]. Similarly, stretch frequency up to 1 Hz induces a contractile phenotype in MSCs and SMCs, whereas static conditions and higher frequencies do not [12-15]. In addition to influencing MSC differentiation, the deposition of ECM factors like elastin and collagen is commensurate with defined mechanobiological stimuli. For example, circumferential MSC conditioning increases elastin content, whereas uniaxial stretch increases collagen content, thereby altering the flexibility or stiffness of the tissue, respectively [16, 17]. Thus, mimicking *in-vivo* strain direction, frequency, and magnitude will result in more accurate *in-vitro* tissue development and is critical for modeling tissues subject to unique biomechanical environments.

The femoropopliteal artery (FPA) is one of the most flexible arteries in the body, accommodating a range of radial, axial, and angular contortions that combine during standing, walking, and crouching movements [18, 19]. Moreover, the FPA’s dynamic biomechanics during limb flexion and its impact on hemodynamics have been implicated in the FPA’s frequent formation of atherosclerotic plaques and occlusive disease progression [18, 20]. Correspondingly, the FPA is also the most likely tissue to be afflicted by peripheral artery disease (PAD), harboring 80% of the 200 million global cases, and disease management treatments exhibit dismal long-term success rates, as low as 39% after five years [21, 22]. A bio-hybrid soft robotic (BSR) bioreactor platform that emulates dynamic FPA biomechanics *in-vitro* will enable researchers to investigate the mechanobiological mechanisms that lead to PAD, access more relevant treatment simulations to improve currently high therapy failure rates, and manufacture tissue engineering vascular grafts (TEVGs) *ex vivo*.

A 3D-printed BSR bioreactor is a promising vessel to mature vascular tissue under patient-, disease-, and lifestyle-specific mechanical stimuli, resulting in personalized tissue shapes and structures that optimally fit and function alongside surrounding vessels. While lab-grown TEVGs have the potential to overcome poor synthetic graft patency rates and alleviate shortages in autograft tissue availability that exclude 40% of patients, there are still many challenges to overcome regarding TEVG durability and mechanical mismatch [23, 24]. Recent venous clinical trials indicate that current TEVGs are prone to spontaneous stenosis due to mechanical mismatches [25]. Previously, TEVGs have been biomanufactured to match the tensile and burst strength of autologous tissue [26-28]. Manufacturing of TEVGs that are suitably compliant for use in environments prone to flexion, such as the FPA, remains elusive. Mimicking native vascular tissue morphology at both the cellular and ECM levels is crucial to growing mechanically robust TEVGs capable of long-term homeostasis [29, 30].

In this project, by developing a first of its kind BSR bioreactor, we demonstrate soft robotics’ potential for conditioning tissues. Using 3D printable materials and a simulation-based design approach, our BSR bioreactors were rapidly developed and personalized, making them befitted for use in patient-specific medicine and research, where no two clinical presentations are the same. We identified Platsil OO-20 as a biocompatible hyperelastomeric silicone suited for both cell culture and PneuNet soft robot fabrication. Characterization of our BSRs actuation modalities revealed that the devices emulate physiologically relevant environmental forces zonally exerted on the FPA, such as angular flexion (AF; 0° to 140°) and radial distension (RD; 0% to 22%), or a combination of angular flexion and radial distension (AR) [19, 31]. Finally, the manufactured BSRs were programmed with angular flexion, radial distension, and angular-radial conditioning regimens and applied to mechanically stimulate MSC monolayers over 24 hours. Analysis of cytoskeletal actin filament orientation revealed that each conditioning regimen induced distinct cellular alignment and high coherency compared to static controls. Similarly, type IV collagen production and phenotypic switching of MSCs to alpha-smooth muscle actin expressing (α-SMA^+^) cells were upregulated in conditioned groups versus static controls. Ultimately, we show that PneuNet soft robots can be used to impart dynamic multi-axial mechanobiological stimuli which induce distinct MSC orientations and smooth muscle cell phenotypes.

## 2. Materials & Methods

### 2.1 Silicone Preparation

Poly(dimethylsiloxane) (PDMS, Dow Corning; Midland, USA) was prepared by combining Sylgard 184 and its curing agent at a 10:1 ratio, respectively, as per the manufacturer’s instructions. PDMS was then degassed in a vacuum chamber for 30 minutes at 0.8 MPa to remove air bubbles created during the mixing process. Platsil OO-20 (Polytek; Easton, USA) was prepared by mixing curing agents A and B at a 1:1 ratio, then degassing for 10 minutes at 0.8 MPa.

### 2.2 Silicone Disk Fabrication

To screen PDMS, Platsil-20, and Visijet M2-Elastomeric Natural (ENT, 3D Systems; Rock Hill, USA) as potential soft robotic cell culture candidates, each material was molded (Platsil-20 & PDMS) or printed (ENT) into 500 μm tall, 6 mm diameter disks. Molds for casting PDMS and Platsil-20 discs were printed from polylactic acid (PLA; Filaform, Adelaide, AU) filament using fused deposition modeling (FDM; Prusa MK3s, Prague, Czech Republic) and a 0.1 mm layer height. After preparation, PDMS and Platsil-20 silicones were cast into the disk molds and cured for 2 hours at 80°C and 60°C, respectively. ENT cell viability disks were printed with a Projet MJP 2500 (3D Systems, Rock Hill, USA) multijet 3D printer. The wax support material was removed via a 1hr steam bath followed by a 1hr heated mineral oil submersion bath. After the oil bath, prints were thoroughly cleaned with hot water and dish soap. In preparation for cell culture, all silicone disk and BSR samples were sterilized by submersion in 70% ethanol for 10 minutes, then exposed to UV light for 20 minutes, and washed with sterile PBS.

### 2.3 Cell Culture

Human adipose-derived mesenchymal stem cells (MSCs, #SCRC-4000; ATCC, Manassas, USA) were used in both viability and BSR cellular conditioning experiments. MSCs were cultured in MSC basal medium (#PCS-500-030; ATCC) mixed with MSC growth kit (#PCS-500-040; ATCC) and 200 µg/ml G418 antibiotic (Thermo Fisher, Seventeen Mile Rocks, Australia). Cells were cultured in a humidified incubator at 37°C and 5% CO2 for cell expansion, viability, and conditioning experiments under QUT human research ethical approval #1900000907.

### 2.4 Cell Attachment and Viability on Silicone Elastomers

PDMS, Platsil-20, and ENT were each subject to a cellular attachment and cytotoxicity study to determine which materials were suitable for bio-hybrid soft robot fabrication. To promote cellular attachment, PDMS, Platsil-20, and ENT silicone elastomer disks were incubated with or without 10 µg/ml fibronectin (#33010-018, Invitrogen; Carlsbad, USA) and suspended in phosphate-buffered saline (PBS) for 24 hours at 37°C. After fibronectin treatment, discs were washed with PBS and transferred to a fresh 48-well plate. MSCs were suspended with each silicone disc at a density of 10,000 cells/cm^2^ for two hours. After two hours, cell-seeded discs were washed with PBS to remove unattached cells, and fresh media was administered. After incubating for 24 hours, cells were live-dead stained with 2 µM calcein acetoxymethyl and 4 µM ethidium homodimer-1 (#L3224, Thermo Fisher) and imaged using fluorescent microscopy to compare cell density and viability between each silicone substrate. Fibronectin-mediated cell attachment and silicone live-dead experiments were carried out in triplicate. Live and dead cells were counted using ImageJ and analyzed with Prism-GraphPad to compare cell adhesion and viability between groups.

### 2.5 Bio-hybrid Soft Robot Design & Simulation

Autodesk Fusion360 (Mill Valley, USA) was used for both computer-aided design (CAD) and finite element analysis (FEA) in a simulation-based iterative design workflow prioritizing angular actuator displacement. Publicly available material properties of Platsil-20 & Ecoflex OO-20 (Smooth-On Inc, Easton, USA) were used to create a profile for a hyperelastic rubber-like material. Fusion360 defines hyperelastic materials as incompressible and fully elastic using the 2-constant standard Mooney-Rivlin model where the distortional constants A01 and A10 define the deformation constant (D1) through the formula D1 == (A10 + A01) × 10^3^.

### 2.6 Bio-hybrid Soft Robot Fabrication

BSRs made of Platsil-20 were cast using a two-part 3D printed mold. Molds for casting BSRs were printed from 3D Tuff resin (Monocure 3D, Sydney, Australia) using a digital light projection printer (DLP; Creality LD-002R, Shenzhen, China). Molds were post-processed after printing with an 8-minute wash in 95% isopropyl alcohol followed by 4-minutes of additional curing under a 405 nm ultraviolet light.

Once prepared, Platsil-20 silicone was poured into top and bottom BSR component molds and placed onto a hotplate at 60°C for at least 2 hours. When solidified, the top and bottom Platsil-20 BSR components were gently separated from their molds and adhered to one another with a thin layer of silicone, then combined and placed back onto the hotplate for an additional 2 hours (Fig 1B).

**Figure 1.**
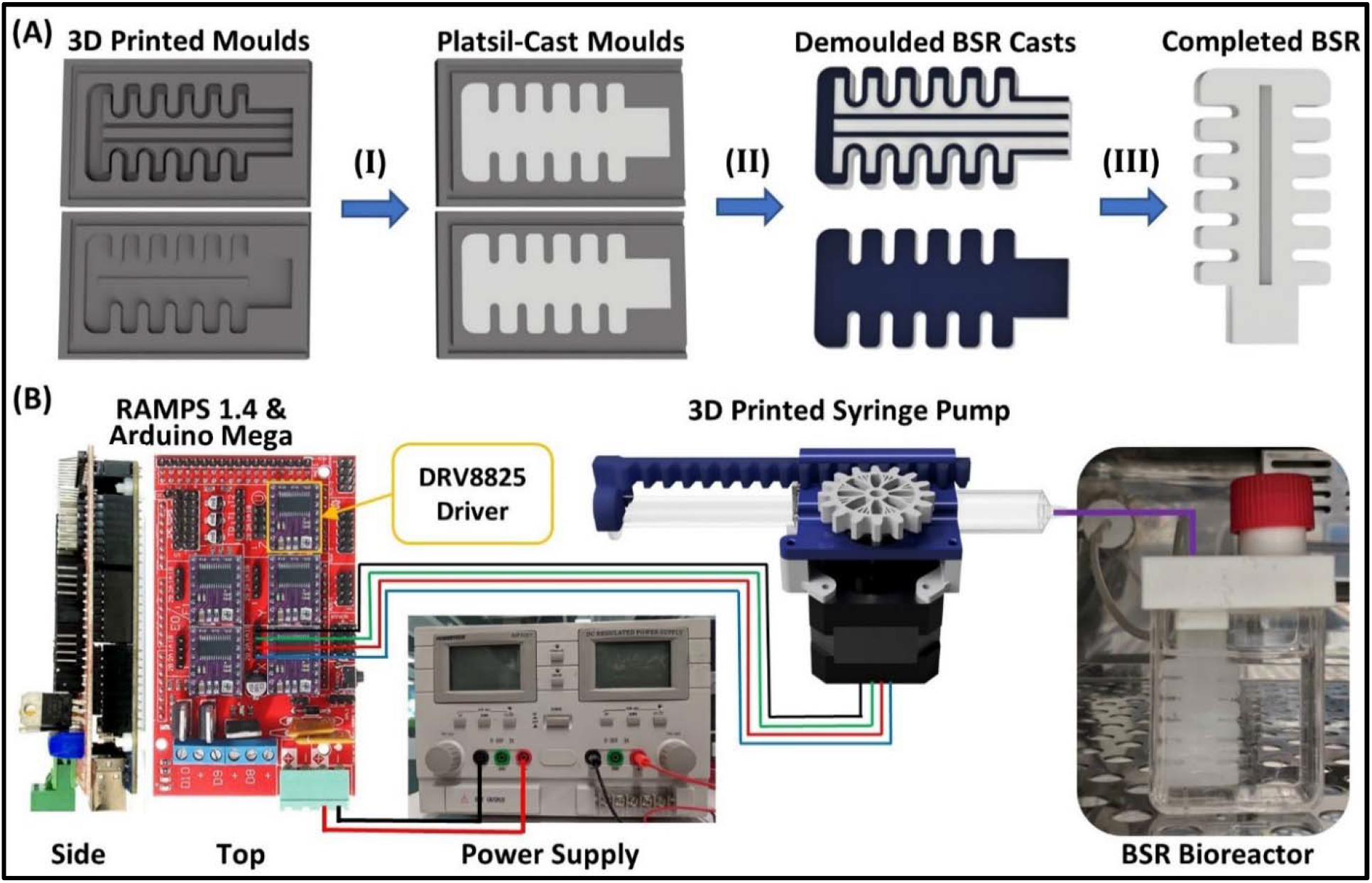
Bio-hybrid soft robots (BSRs) are fabricated using 3D printed molds and computer-controlled through pneumatic flexion from custom syringe pumps. Part A outlines a step-by-step process for manufacturing BSRs from Platsil-20 (white) using 3D printed molds (gray). (I) Platsil-20 parts A & B are mixed, degassed, and poured in the BSR molds. (II) The top and bottom BSR components are cured on a hotplate at 60°C for 2 hours and removed from the molds. (III) Each component’s inner face is then pressed together and placed on the hotplate to be fused, resulting in a completed BSR. Part B illustrates the core components of the BSR control unit and cell culture apparatus. The control unit comprises an Arduino Mega 2560 microcontroller fitted with a Ramps 1.4 shield and DRV8825 stepper drivers used to control a 3D printed syringe pump. A custom cell culture apparatus was made from a T25 culture flask modified with a 3D printed lid incorporating BSR and filtered cap attachment ports.

### 2.7 Imaging of DLP-printed moulds and resultant silicone structures

Scanning electron microscopy was performed on DLP resin printed moulds used for creating the BSRs. DLP-printed moulds were sputter-coated with 4 nm platinum (Leica EM ACE600, Wetzlar, Germany) And imaged with a Phenom XL G2 Desktop SEM (Thermo Fisher Scientific, Thermo Fisher, Waltham, Massachusetts) at a 6mm working distance at 5 kV.

To visualise the microgrooves created via the moulding process, DLP-printed resin moulds and cast silicone BSRs were visualised under a Nikon SMZ25 stereo zoom microscope (Nikon, Tokyo, Japan) with a 1x objective at 2.5x and 7x zoom and captured with a Nikon DS-R12 (Nikon, Tokyo, Japan) 16.25 megapixel mounted camera.

### 2.8 Bio-Hybrid Soft Robot Control

BSR actuators were pressurized by rack-and-pinion syringe pump that was FDM 3D printed from acrylonitrile butadiene styrene (ABS; Filaform, Adelaide, AU) filament. The pump was printed from ABS to prevent warping due to high temperatures generated by the stepper motor from continuous use. The syringe pump was controlled by an Arduino MEGA 2560 (Ivrea, Italy) and a RAMPS 1.4 shield (Maker Store, Melbourne, Australia) fitted with DRV8825 stepper drivers (Maker Store, Melbourne, Australia) (Fig 1B). Initial prototypes printed from PLA did not maintain structural integrity over long periods and warped due to the stepper motor heating above the glass transition temperature of PLA (60°C). However, due to its higher glass transition temperature (105°C), components printed from ABS functioned for back-to-back 24-hour sessions without overheating and failure.

To house the BSRs, a custom culture chamber was made from a modified T25 flask, and FDM 3D printed thermoplastic polyurethane (TPU; Ninjatek, Manheim, USA) lid (Fig 1B). The lid incorporates a port that is compatible with a T25 filter cap for media changes and to facilitate sterile air diffusion. In addition, an extended pneumatic port built into the lid ensures that the BSR remains in place over long periods of flexion.

### 2.9 Bio-hybrid Soft Robot Cell Culture & Staining

BSRs were coated with 10 µg/ml of fibronectin for 24 hours prior to seeding with MSCs. In addition, a recessed region on the surface of the BSR was created to improve fibronectin & MSC coating/seeding accuracy. The recessed region possessed a surface area of 0.6 cm^2^ and was seeded with a density of 4×10^4^ MSCs/cm^2^ suspended in 300 µl of media for 2 hours. After 2 hours, seeded BSRs were washed with PBS to remove unattached cells and placed into a custom culture flask, and expanded for 3 days. After 3 days, cells on RD, AF, and AR devices were conditioned for 24 hours (Table 1). Cells undergoing RD conditioning were subject to a 13% stretch cyclically at 0.67 Hz. Cells conditioned with AF were subject to a 153° angle of actuation cyclically at 0.5 Hz. Static control (SC) and tissue culture plastic (TCP) controls underwent identical media change and material processing steps as the conditioned groups but remained static and were not actuated.

**Table 1:** Bio-hybrid soft robot culture conditions: bending angle, percent radial distension, and rate of actuation (Hz), with 3 independent cultures for each BSR condition.

| BSR Group | Banding Angle / Frequency (Hz) | Percent Radial Distension / Frequency (Hz) |
| --- | --- | --- |
| Static Control (SC) | 0° / 0.0 Hz | 0% / 0.0 Hz |
| Angular Flexion (AF) | 153° / 0.5 Hz | 0% / 0.0 Hz |
| Radial Distension (RD) | 0° / 0.0 Hz | 13% / 0.67 Hz |
| Angular + Radial (AR) | 153° / 0.5 Hz | 13% / 0.67 Hz |

After culture, cells were fixed with 4% paraformaldehyde (PFA) for 20 minutes at room temperature and washed with PBS (3x). Before adding primary antibodies, fixed cells were permeabilized with 0.1% Triton X-100 for 10 minutes, then washed with PBS (2x) and incubated in PBS + 3% fetal bovine serum (FBS) for 20 minutes at room temperature. A primary antibody incubation of rabbit anti-human Col IV (1:1000; AbCam, ab6586, Cambridge, United Kingdom) and goat anti-human α-SMA (1:1000; AbCam, ab21027) in PBS + 1% FBS was applied for 18 hours. Cells were then washed in PBS and treated with a secondary antibody incubation containing donkey anti-rabbit AlexaFluor 555 (1:500; Invitrogen), donkey anti-goat AlexaFluor 647 (1:1000; Invitrogen), and phalloidin AlexaFluor 488 (5 U/ml; Invitrogen) in PBS + 3% FBS for 2 hours. Cells were once again washed with PBS (3x) and counterstained with DAPI (1:1000) for 5 minutes at room temperature. After a final series of PBS washes (3x), cells were imaged using confocal microscopy (Nikon A1R, Tokyo, Japan).

### 2.10 Image Analysis

The orientation and distribution of MSC cytoskeletal actin filament were analyzed using the ImageJ plugin, OrientationJ [32]. Images of phalloidin-stained actin filaments were separated from merged z-stacks and analyzed with a cubic spline gradient and local α window of 2 pixels. Type IV collagen and α-SMA expression between conditioning regimens was analyzed by comparing normalized fluorescent intensity. Min-max normalization (I-I_min_)/(I_max_-I_min_) was used to rescale intensity values between 0 and 1, enabling comparison of replicates imaged with alternate microscope settings, as performed previously [33]. The particle analysis tool in ImageJ was used to count DAPI-stained nuclei for cell density quantification.

### 2.11 Statistical Analysis

Differences between cell viability (n = 4), BSR angular flexion (n = 3) and radial distension (n = 6) characterization, nuclear density (n = 3), and immunofluorescence (n = 3) experimental groups were evaluated via unpaired t-test and statistical significance was defined as P < 0.05.

## 3. Results & Discussion

### 3.1 Elastomer Characterization

Poly(dimethylsiloxane) (PDMS) is the most common elastomer in microfluidic cell culture applications due to its cytocompatibility and optical clarity. A range of mechanobiological devices already employ PDMS as a stretchable cell culture substrate [35-37]. However, PDMS is not ideal for soft robotics applications due to a poor elongation at break (∼140%) [38]. In addition to PDMS, we identified Platsil-20 and ENT as potential candidates for their ideal mechanical properties and ease of manufacture. Platsil-20 has a shore hardness of OO-20 and elongation at break of 964%, but Platsil-20 becomes opaque when cured, preventing live cell imaging in 3D culture platforms. Alternatively, ENT is an elastomeric resin from 3D systems which can be directly 3D printed using a Projet MJP 2500 MultiJet Printer. Being a 3D printable elastomer, ENT could allow for the fabrication of more complex and microscale PneuNets. Moreover, anatomical surface geometries for highly relevant mechanobiological stimuli would be much easier to include in device design and fabrication workflows. Unfortunately, the elongation before break of ENT is much lower than Platsil-20 at 160-230%, and ENT’s optical clarity is also not suitable for imaging in a 3D culture format.

Platsil-20 and ENT have not been previously documented for use in cell culture. Therefore, we characterized the biocompatibility of these materials through the attachment and viable growth of MSCs. MSCs are relevant to vascular tissue physiology and remodeling because they can differentiate into multiple somatic cell types, such as smooth muscle cells (SMCs), which are critical for maintaining vascular tone and generating collagen IV and elastin ECM components [17, 30, 39]. To assess their potential, we conducted a series of cell viability and attachment assays in which MSCs were seeded onto 6mm discs of each material with and without fibronectin coating. Positive controls included cells cultured on well plates with and without fibronectin coating. After a 24-hour incubation period, each group was live-dead stained and imaged (Fig 2).

**Figure 2.**
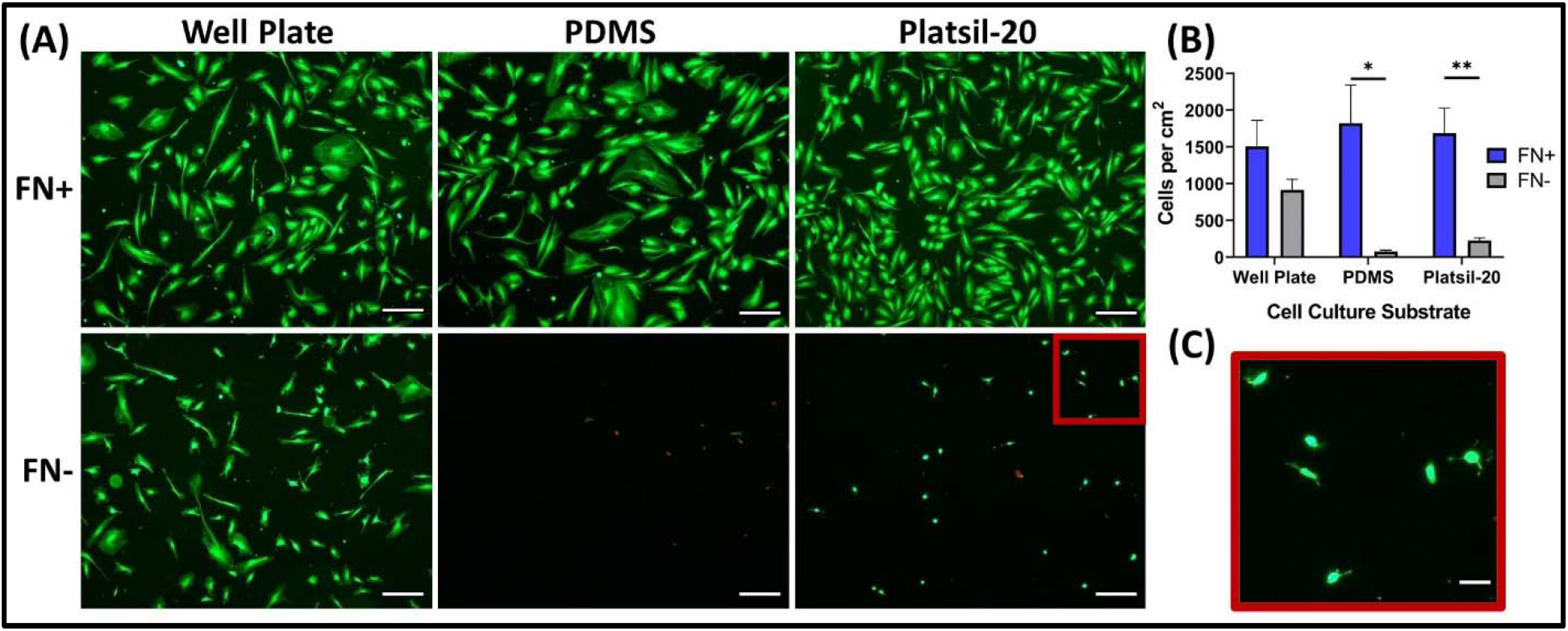
Fibronectin treated silicone supports MSC culture. (A) Representative images of live-dead stained MSCs cultured on the surface of either a 48-well plate, 6 mm PDMS disc, or 6 mm Platsil-20 disc coated with or without fibronectin (scale bar = 100 μm). (B) A bar graph showing the number of live cells counted on each surface with (blue) and without fibronectin (grey) (n = 4). P values were obtained via an unpaired t-test. (C) A magnified region of MSCs lightly attached to and spreading on Platsil-20 without fibronectin (scale bar = 25 μm).

When coated with fibronectin, MSCs attached to the tissue culture plates, Platsil-20, and PDMS with equal efficiency (Fig 2A, 2B). However, without fibronectin coating, fewer cells were attached to the tissue culture plate, and virtually no cells were attached to Platsil-20 or PDMS (Fig 2A, 2B). Cellular attachment to Platsil-20 without fibronectin appeared very weak, as the cells are mostly rounded with slight body elongation (Fig 2C). Both PDMS and Platsil-20 possessed the same cytocompatibility as the well plate control with fibronectin, producing adhered, confluent monolayers comprised of cell viabilities near 100%. Conversely, ENT was cytotoxic, with no live cells attached to the material’s surface or on the well plate adjacent to the silicone disc (Fig S1). Moreover, ENT was noticeably autofluorescent, making cell imaging difficult. Altogether, fibronectin-coated Platsil-20 possessed a suitable cellular attachment and cytocompatibility to be used as a cell culture substrate in BSR experiments.

### 3.2 Bio-hybrid Soft Robot Design & Characterization

Previous biomechanical investigations into the effects of limb flexion on FPA tortuosity have highlighted angular flexion (AF), cyclic radial distension (RD), torsion, and axial stretch as the four primary modes of mechanical stimuli on arterial tissue [19, 20, 40]. We aimed to include as many of these modalities into a single device as possible using a simulation-based iterative strategy (Fig 3A). Published soft pneumatic actuator designs were assessed for their ability to emulate key vascular stress modalities, ease of fabrication, and practical cell culture functionality [1, 4, 41, 42]. Ultimately, a multichannel pillared PneuNet design was selected for its ability to emulate popliteal AF (Fig 3B & 3C). Moreover, axial stretch can be achieved in our device if both PneuNets are pressurized simultaneously. A third modality was incorporated by placing a hollow space between the two pillared PneuNets that, when pressurized, expands to create RD (Fig 3B & 3D). Future advancements in soft robotic design and fabrication, such as rapid fiber-reinforced actuators, will broaden the biomimicry applications of BSR bioreactors to new tissue and organ systems [43]. Additionally, improvements in BSR miniaturization will increase the feasibility of high throughput in-vitro models by reducing experimental material costs.

**Figure 3.**
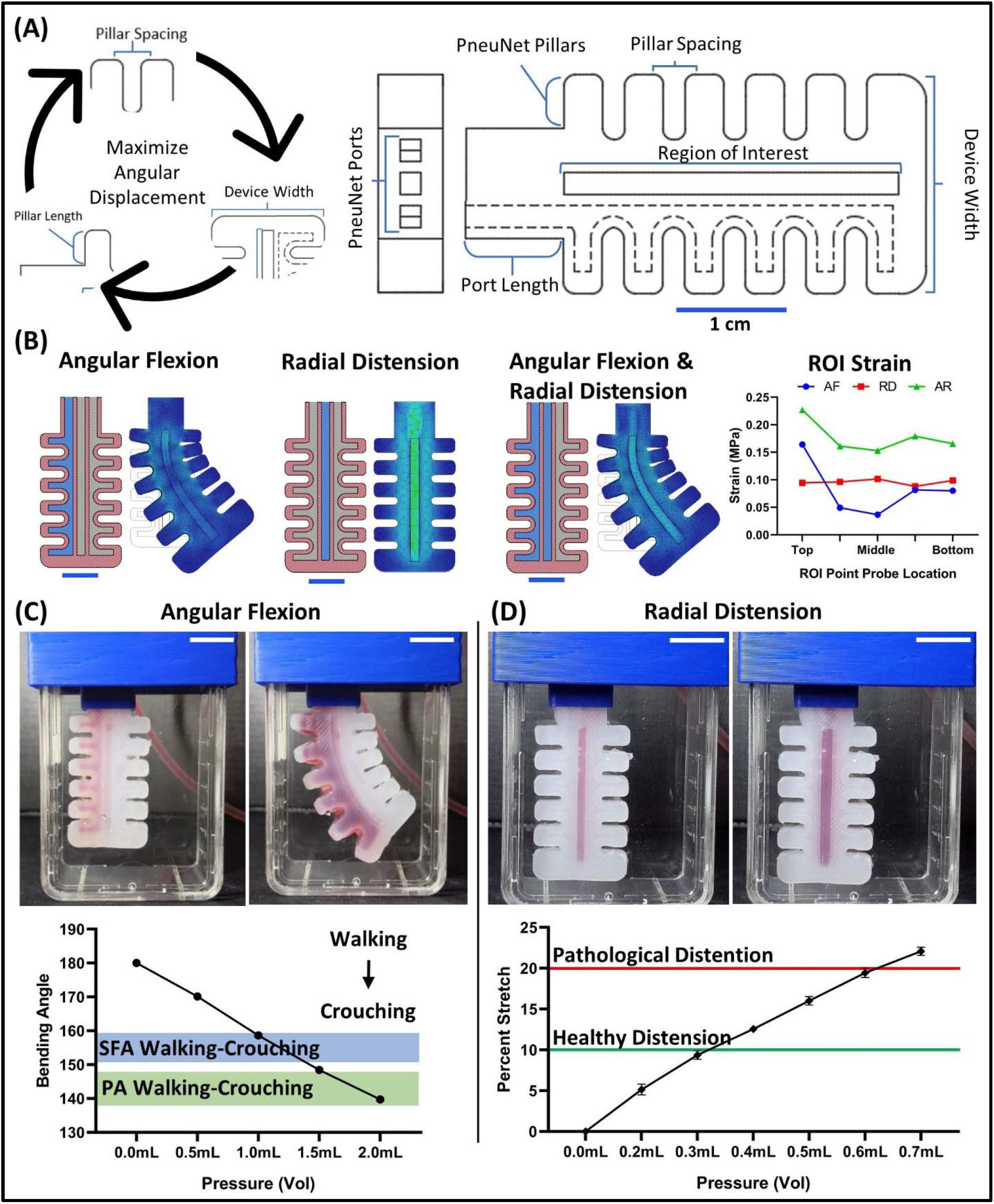
BSRs were designed and optimized using a simulation-based iterative workflow that enables multi-axial actuation to emulate femoropopliteal physiology. (A) Geometric and design parameters, including pillar length, pillar spacing, and device and PneuNet width were optimized to achieve the highest potential BSR angular displacement. The final design exhibited a reduced volume and recessed ROI to concentrate cellular attachment. (B) Angular flexion (AF), radial distension (RD), and combined (AR) actuation modes were simulated and analyzed with five evenly spaced simulation ROIs to assess relative differences in strain. The PneuNet channel pressurized to achieve each respective actuation mode is highlighted in blue. (C) A linear relationship between the volume of injected air and the bending angle was found and used to design later conditioning studies. AF of the BSR ranges between 180° to 140.6° from PneuNet pressurization of 0 to 2 ml of air, respectively (n = 3). The range of bending angles across the SFA between walking, sitting, and crouching movements are highlighted in blue. Similarly, the same range of PA bending angles are highlighted in green. (D) We show that healthy (10%) and pathological (20%) RD can be achieved by pressurizing the device with 0.3 ml to 0.7 ml of air (n = 6). All scale bars = 1 cm.

A simulation-based iterative workflow was utilized to optimize BSR design for maximum angular displacement while constraining total BSR volume to fit within a T25 culture flask. We found that pillar spacing, minimal radial PneuNet width, and high PneuNet length to width ratio had the greatest effect on maximizing angular displacement. Next, we extruded a recessed channel over the surface of the BSR to better control fibronectin coating and cell seeding on a hydrophilic surface. By employing a simulation-based iterative strategy, the silicone required to fabricate each BSR was reduced from 5.5 ml to 3.1 ml, resulting in a 43% BSR volume reduction compared to initial prototypes. FEA simulations of AF, RD and AR actuation were used to gain insight into differences in mechanical strain across the BSR region of interest (ROI) for each conditioning modality. Strain measurements for each mode were acquired from five pointprobes spaced evenly across the BSR ROI and graphed in Figure 3B. The AF and RD modes simulated a flexion of 153° and 10% radial stretch, respectively, whereas the AR mode simulated a simultaneous combination of AF and RD. We found that strain remained consistent across the RD ROIs but fluctuated across the AF ROIs. The AR simulation indicated the highest overall strain with but with less fluctuation between point probes than the AF mode. While the combined AR simulations applied angular and radial actuation simultaneously, cyclic radial and angular actuations are not synchronised in human physiology nor in our experiments. Still, these simulations provide insight showing that AF creates a spectrum of strain that will produce unique mechanobiological stimuli on the conditioned cells. Asynchronous cyclic simulations could provide future insight into how multi-axial strain directions and magnitudes affect heterogeneous tissue maturation.

To design conditioning regimens for BSR cell culture experiments, we first characterized physiological bending angles from actuating the AF PneuNet and percent radial stretch from the RD PneuNet (Fig 3C & 3D). The bending angle was characterized by pressurizing one of the peripheral PneuNets with various volumes of air and measuring the resultant angle formed between three centerline points evenly spaced across the top, middle, and bottom of the BSR ROI. A line was digitally drawn between the top and middle points and the bottom and middle points using ImageJ. Then, the bending angle was calculated as the angle formed at the intersection of each line on the middle point. By pressurizing devices with 0 to 2 ml of air, we achieved a bending angle of 180° to 140°, respectively (Fig 3C). Previous literature identifies acute bending angles across the FPA during walking and sitting limb flexion ranging between 141° and 158° [19]. Thus, our results indicate that the BSRs are capable of emulating acute FPA bends during knee flexion. In addition, RD under various applied air pressures was determined by measuring the change in radial width of dots across the BSR ROI. Our results show that pressurizing the RD PneuNet with 0 ml to 0.7 ml of air induced a 0% to 22% RD (Fig 3D). FPA RD of 10% is typical during a normal cardiac cycle for healthy individuals, whereas pathologies such as hypertension can result in distensions exceeding 20% [31]. Therefore, the BSRs can apply both healthy and pathological stimuli to cells by pressurization the center PneuNet with 0.3 ml to 0.7 ml of air.

### 3.3 Cytoskeletal Orientation & Distribution

Figure 4 shows immunofluorescent z-stack confocal images of MSCs cultured on SC, AF, RD, and AR BSR conditioning regimens. All samples were seeded with 4×10^4^ MSCs/cm^2^, expanded for 3 days, then subject to their perspective conditioning regimen for 24 hours. The functional alignment of cells can be precisely measured by analyzing their cytoskeletal orientation. Here we used the OrientationJ tool to visualize and quantify the orientation and distribution of cytoskeletal actin filaments (phalloidin staining) for each BSR conditioning regimen [32]. The actin filaments are digitally colored based on their alignment, with those aligned in a similar direction being visible in a similar color (Fig 4D). The discordant color pattern in the SC image of Figure 4D shows that the cytoskeleton of statically cultured MSCs is disordered and does not cohesively align in one direction. In comparison, images of the AF, RD, and AR conditioned MSCs show that each regimen induces a unique cytoskeletal orientation. While RD conditioned MSCs aligned longitudinally to the ROI, AF conditioned MSCs have a circumferential cytoskeletal alignment that mimics what would traditionally be expected by SMCs within the arterial tunica media [30, 39, 44]. Furthermore, MSCs from the AR conditioning regimen possess a diagonal cytoskeletal alignment within the ROI, showing that multi-axial AF and RD actuation can be combined to produce tight distributions of any desired cell orientation.

**Figure 4.**
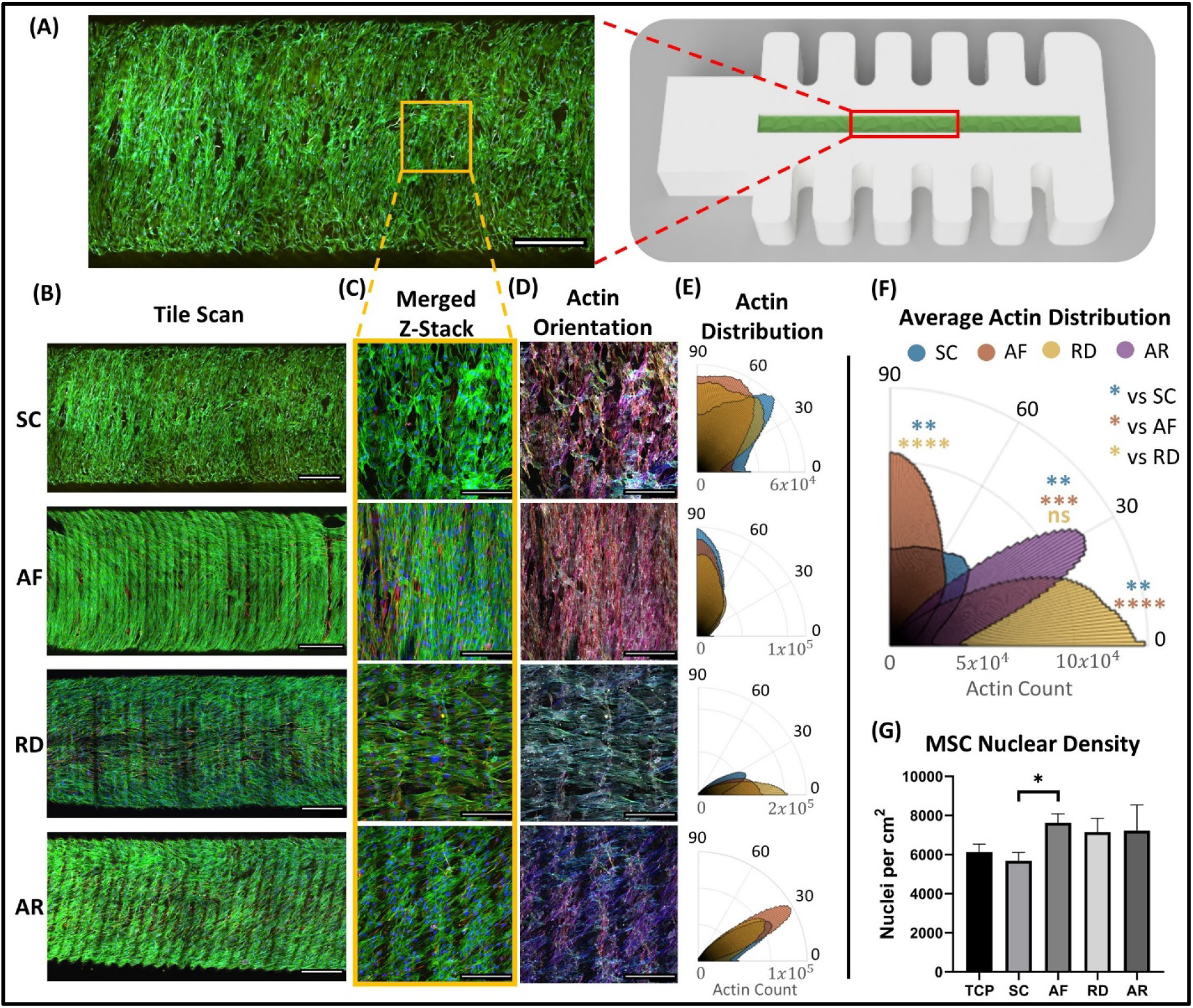
BSR conditioning affects MSC cytoskeletal orientation after only 24 hours. (A) A 3D rendering showing that tile scans, and focused 10x and 20x images used for actin analysis were acquired from the centerline of the BSR ROI (scale bar = 500 µm). (B) Tile scan z-stacks of MSCs cultured on BSRs conditioned by static control (SC), angular flexion (AF), radial distension (RD), and angular with radial (AR) conditioning types. The z-stacks merge four channels to show actin filament orientation (green), cell nuclei (blue), type IV collagen (red) and α-SMA (yellow) (scale bars = 500 µm). (C) Representative merged 20x z-stacks of MSCs cultured in each conditioning regimen (scale bars = 100 µm). (D) Orientation analysis of actin filament for each conditioning type in which the actin filament have been digitally stained corresponding to filament orientation (scale bars = 100 µm). (E) Distribution analysis of actin filament from 10x images for each conditioning type (n = 3). (F) A graph showing the average actin filament distribution and significance comparisons between each condition type. (G) Nuclear density of MSC cultured in each conditioning type (n = 3).

Actin filament distribution data in Figure 4E shows that each BSR conditioning regimen induced highly ordered cytoskeletal alignment. Moreover, the mean actin distribution angle was significantly different between the SC and all conditioned groups, except for the RD to AR comparison (Fig 4F). As expected, the SC group displayed the highest level of disorder, with a mean actin orientation of 65.8° and a standard deviation of actin filament orientation of 33.4°. We also observed that the actin filament of AF, RD, and AR groups were oriented with unique dominant alignments of 87.7°, 13.6°, and 32.4°, respectively. Interestingly, the level of disorder between conditioned groups decreased progressively, with the mean standard deviation of actin filament orientation decreasing from 29.6° to 20.2° between AF and RD regimens. Ultimately, the AR conditioning regimen resulted in the lowest disorder, with a mean actin filament deviation of 17.7°.

Longitudinal actin alignment in the RD regimen corroborates previous observations that cells will align perpendicularly to the direction of uniaxial stretch [13, 14]. Therefore, the circumferentially aligned cells from the AF regimen could have instead been expected from cyclic axial stretch, and perhaps suggests the BSR’s angular component did not distinctly augment cell alignment versus linear extension alone. Recent literature shows that SMC orientation in the tunica media does not express a perfect circumferential alignment but orients in a helical or spiral-like pattern around a vessel that varies depending on the vessel being studied [45]. Thus, the diagonal cell orientation observed from AR conditioning could be leveraged to mimic these helical structures more closely than uniaxial stretch. Moreover, while each BSR’s cell orientation ROIs were selected from the flat center of its fibronectin-coated recess, the curved and vertical sides of the recesses contained more horizontal cell orientations (Fig 3B). Therefore, the combination of mechanical stresses with various surface topologies may enable the fabrication of more complex 3D tissue structures in future studies (Fig S2). The additively manufactured BSR molds resulted in microscale 3D printing layer lines across the BSR and its recesses and imaged ROIs. Previous studies have shown that similar microscale topological features can affect cellular morphology and alignment [46, 47]. However, no significant cell alignment due to these layer lines was observed for static culture (SC) replicates. Future investigations into how cells respond to competing mechanobiological stimuli would aid decision-making when choosing the most effective methods for guiding tissue maturation.

A similar density of MSCs was found between AF, RD, and AR conditioning groups by counting the number of nuclei present in each image (Fig 4G). While conditioning groups had greater nuclear densities than static and tissue culture plastic control groups, only the AF group had significantly greater nuclear densities than the SC group (P < 0.05). Cells within the SC group appear to have a spread-out synthetic morphology, while cells from the conditioned groups were more elongated and spindle-like. The higher density of nuclei in the conditioned regimens could result from elongated morphologies that enable cells to pack tightly together. Similar observations have been reported in other cell stretching studies, in which mechanically conditioned tissue monolayers contained elongated morphologies and greater numbers of cells [48]. To clearly delineate effects of different condition regimes on cell proliferation, mechanical stimulation experiments should be performed on cell monolayers for longer periods of time.

Closer analysis of cellular morphology across new regions of the BSR ROI, and employment of alternate conditioning regimens is needed to determine whether stimulating MSCs with AF is identical to a uniaxial stretch. Regardless, these results indicate that the BSRs create multimodal conditioning regimens that induce unique MSC alignment, highlighting their potential for disease modeling and tissue engineering. Moving forward, comparisons of cytoskeletal alignment at unique stress points across the AF ROI could yield findings on how tissues develop in response to gradients of stress, such as those found within the popliteal artery [49]. Moreover, alternative regimens could be created to investigate the effects of different stretch magnitudes, frequencies, and rest periods.

### 3.4 Collagen IV & α-SMA Production

To further investigate the effects of the BSR conditioning on MSC function and differentiation, we assessed α-SMA and type IV collagen production after either 4 days of static culture (TCP and SC control groups) or 3 days of static culture followed by 24 hours of conditioning (AF, RD, and AR groups) using immunofluorescent staining. Increased α-SMA production indicates that MSCs are progressing towards a contractile phenotype commonly expressed by smooth muscle cells (SMCs) [50]. Similarly, the production of ECM factors, like type IV collagen after 24 hours of cyclic stretch, further demonstrates MSC phenotypic switching towards SMCs [17, 50, 51]. While differentiating MSCs into SMCs using growth factors like TGF-B is well documented, the use of mechanical stimuli alone to promote SMC differentiation has also been described [52]. Given the high cost of growth factors, vascular tissue engineering methods that employ mechanobiological tools could provide efficient bioreactor platforms for aligned MSC-derived SMC differentiation and vascular tissue manufacturing.

Indeed, our results showed a distinct increase in both type IV collagen and α-SMA^+^ production between mechanically-conditioned and static control groups after only 24 hours of BSR conditioning (Fig 5A & 5B). Furthermore, all conditioned groups expressed significantly more type IV collagen than both static control groups. However, when comparing α-SMA, only the AF group had a significantly higher pixel intensity than static control groups. These data indicate that MSCs begin shifting towards an SMC phenotype after 24 hours of BSR conditioning for all actuation types. Counterintuitively, this data also indicates that multi-axial AR conditioning has the lowest collagen IV and α-SMA expression of all mechanical conditioning groups. Previous research investigating MSC mechanobiology has shown that cells conditioned with equiaxial stress produce less α-SMA than cells conditioned with uniaxial stress, which could explain this behavior [9]. Altogether, these data suggest that the unique mechanobiology within vascular structures located in high flexion or uniaxial stretching regions could induce altered cellular function. Future studies will investigate aspects of BSR tissue maturation over extended periods where we expect to see a more exacerbated difference between cells conditioned with different types of stress. For example, a reduced expression of stiff matrix proteins from equiaxial stress, such as collagen IV, would result in less rigid arterial structures that may be more suitable in environments like the popliteal artery. Moreover, widening the scope of ECM protein analysis could show increased elastin content in AR conditions compared to stiff AF and RD tissues with elevated collagen content. Finally, extended conditioning regimens will likely further differentiate MSCs towards vascular SMC phenotypes but also reduce cellular proliferation compared to the static control groups.

**Figure 5.**
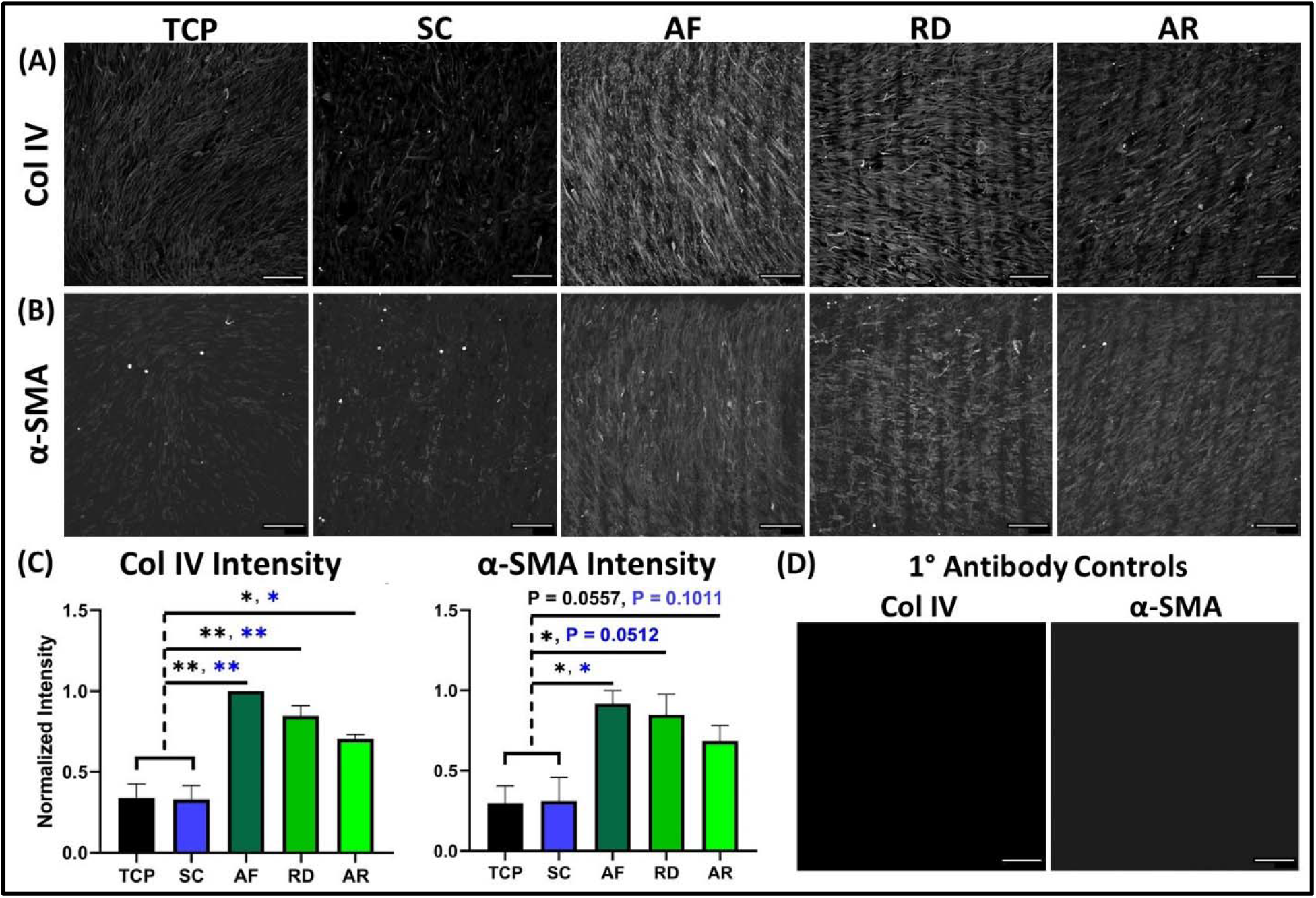
24 hours of BSR conditioning increases collagen IV and α-SMA production in MSCs. (A) Representative grayscale images showing type IV collagen and (B) α-SMA presence between control and test groups after 24 hours of conditioning. (C) Graphs comparing type IV collagen and α-SMA intensity between BSR control and conditioning regimens. Type IV collagen intensities for tissue culture plastic (TCP) and static control (SC) cultures were significantly different from mechanically-conditioned groups, while α-SMA intensity of the TCP control differed significantly from AF and RD groups, but not the AR group. (D) Type IV collagen and α-SMA primary antibody control. All scale bars = 200 µm, n = 3.

## 4. Conclusion

In this investigation, we developed a novel approach for mechanically conditioning cells that leverages bio-hybrid soft robotics (BSR) to induce MSC differentiation and alignment. First, we showed that Plastsil OO-20, a hyperelastomer ideal for soft robotics fabrication, is biocompatible and supports strong cell adhesion when coated with fibronectin. Then we described the methods by which BSRs can be fabricated using widely available 3D printing tools and materials. By characterizing BSR actuation, we showed that our devices emulate bending angles experienced in the FPA during daily limb flexion, from 140° to 180°. We also showed that the same BSR recreates healthy and pathological arterial radial distension (RD) by pressurizing a central PneuNet located below the ROI. Finally, we carried out a cell culture study comparing the effects of typical cell culture against BSRs exerting static, RD, angular flexion (AF), and a combination of angular flexion and radial distension (AR) conditions on MSC cytoskeletal alignment, type IV collage production, and α-SMA production. Orientation and distribution analysis of MSC cytoskeleton revealed that conditioning regimes could be combined to align cells into any desired tissue orientation, with the AR group possessing the tightest alignment. Interestingly, the production of type IV collagen and α-SMA among the mechanically-conditioned groups was highest after AF and lowest after a multi-axial AR actuation regime.

This study is the first application of a hyperelastomer substrate in a bio-hybrid soft robotic bioreactor that directs highly ordered organization and phenotypic switching of MSCs by combining a series of pneumatic networks for multi-axial actuation using advanced manufacturing techniques. While this BSR may replicate aspects of femoropopliteal mechanics, it does not form a cylindrical or patient-specific vessel structure, which can be designed and cast from personalized 3D printed molds in future iterations. Future BSRs will integrate a hollow vessel model and medium perfusion to create a shear stress modality and perform long-term culture studies for biomanufactured vascular grafts and *in vitro* testbeds for medical and surgical treatments on patient-specific femoropopliteal arteries, disease modeling, biomechanical testing of endovascular and medical therapies, and the biomanufacturing of tissue-engineered vascular grafts.

## Acknowledgments

The author would like to thank Mitchell Johnson for assistance with programming the pneumatic control unit, Sabrina Schoenborn and Ryan Daley for productive additive manufacturing and experimental design discussions, A/Prof Dr. Yi-Chin Toh for providing PDMS and silicone tubing, and the QUT Central Analytical Research Facility for assistance with microscopy. This research was supported by an Advance Queensland Fellowship (AQIRF1312018) and fruitful discussions with Dr. Robyn Stokes, Dr. Geremy Farr-Wharton, and Bionics Queensland.

## Supplementary

**Figure S1.**
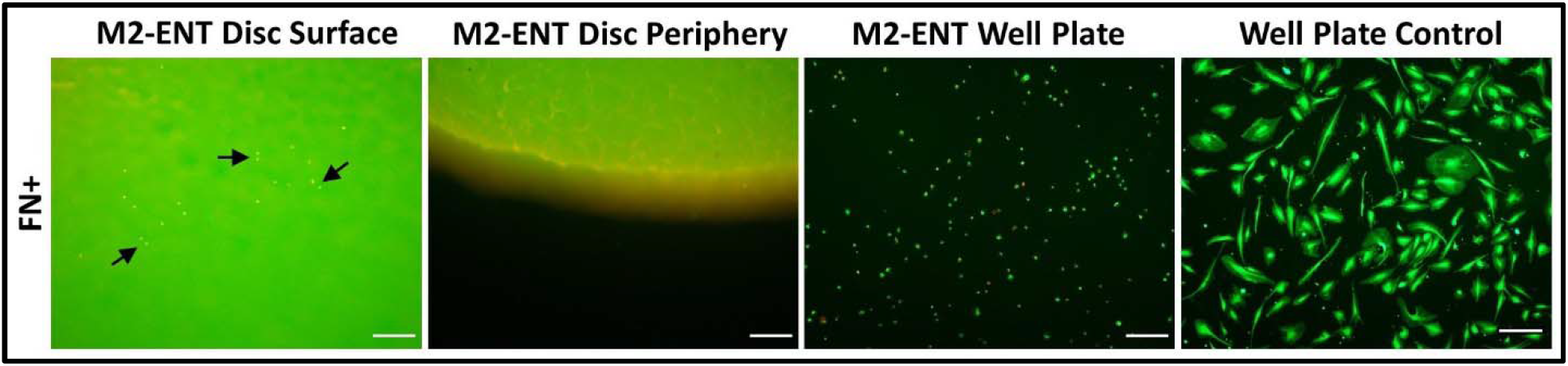
M2 ENT is cytotoxic towards MSCs. ENT is autofluorescent, as well as cytotoxic to MSCs directly on its surface and in the adjacent well-plate. Arrows on the first image point to dead cell clusters resting on the surface of the ENT disc. The final image in the ENT series shows dead MSCs resting adjacent to the ENT disk, at the bottom of the well-plate. Scale bars = 100 μm.

**Figure S2.**
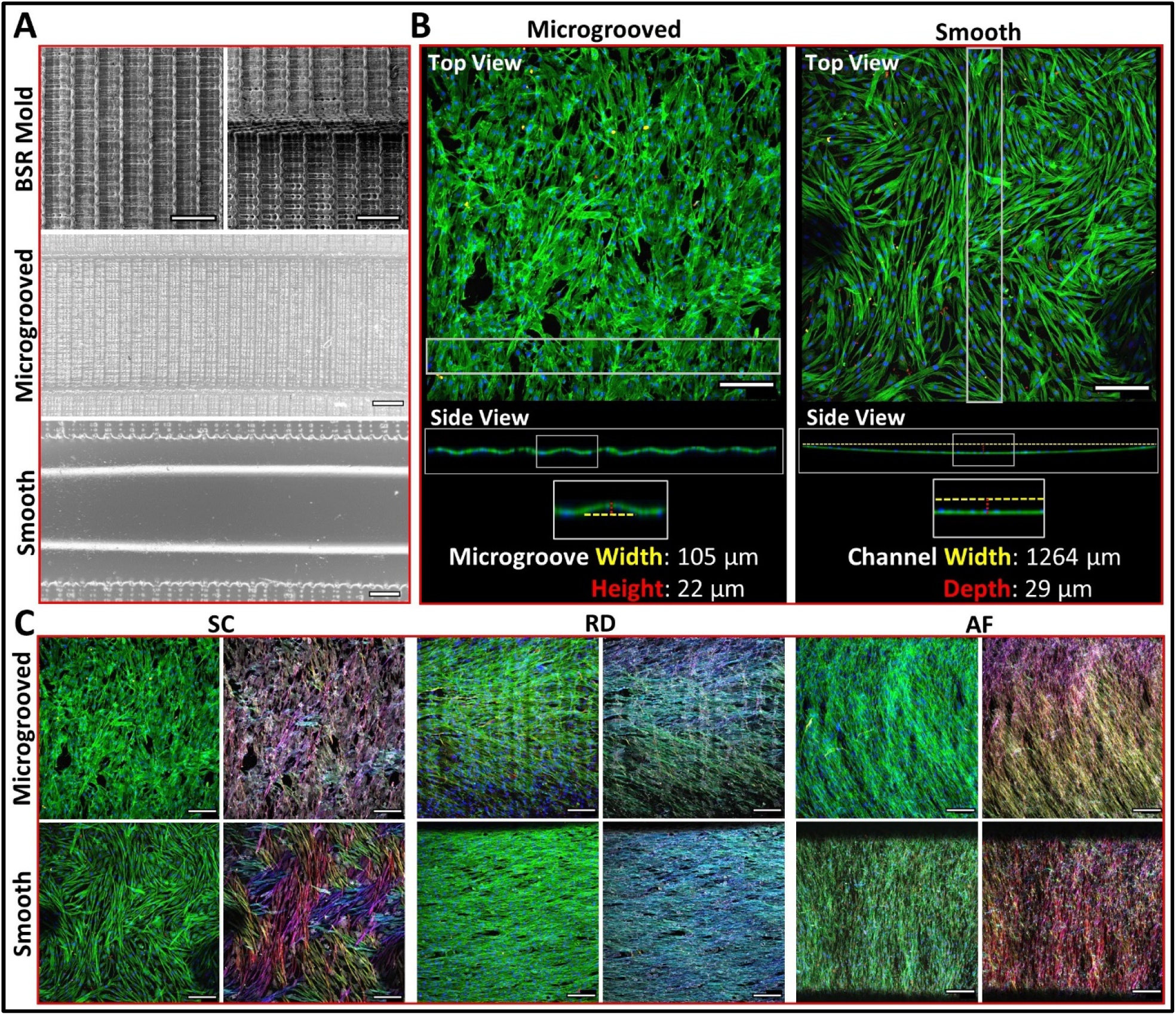
3D printed microgroove topology influences cytoskeletal morphology. (A) Scanning electron microscopy of the microgrooved resin BSR mold and stereo zoom microscopy of the microgrooved and smoothed silicone BSRs images show that unmodified BSRs recapitulate the layer line topology of the 3D printed molds from which they are cast, creating a series of rough microgrooves perpendicular to the ROI. Modified BSRs with a smooth surface were made to determine if 3D printed microgroove topology affects MSC cell culture (scale bars = 300 µm). (B) Here, confocal images show top and side profile views of statically cultured MSCs on microgrooved and smooth BSRs. The side profile views show that MSCs adhere to and match the shape of each perspective surface topology, while the top view shows that MSCs statically cultured on smooth BSRs (n = 1) have larger and more elongated morphologies than the microgrooved BSRs (scale bars = 200 µm). (C) Visually comparing the cytoskeletal morphology of MSCs cultured in static (SC) and mechanical conditions (RD and AF) show that morphological differences are less pronounced between the microgrooved and smooth (n = 1) BSRs than statically cultured samples. However, the actin orientation of MSCs in smooth RD and AF groups appears to be more uniformly aligned than their microgrooved counterparts (scale bars = 200 µm).

**Figure S3.**
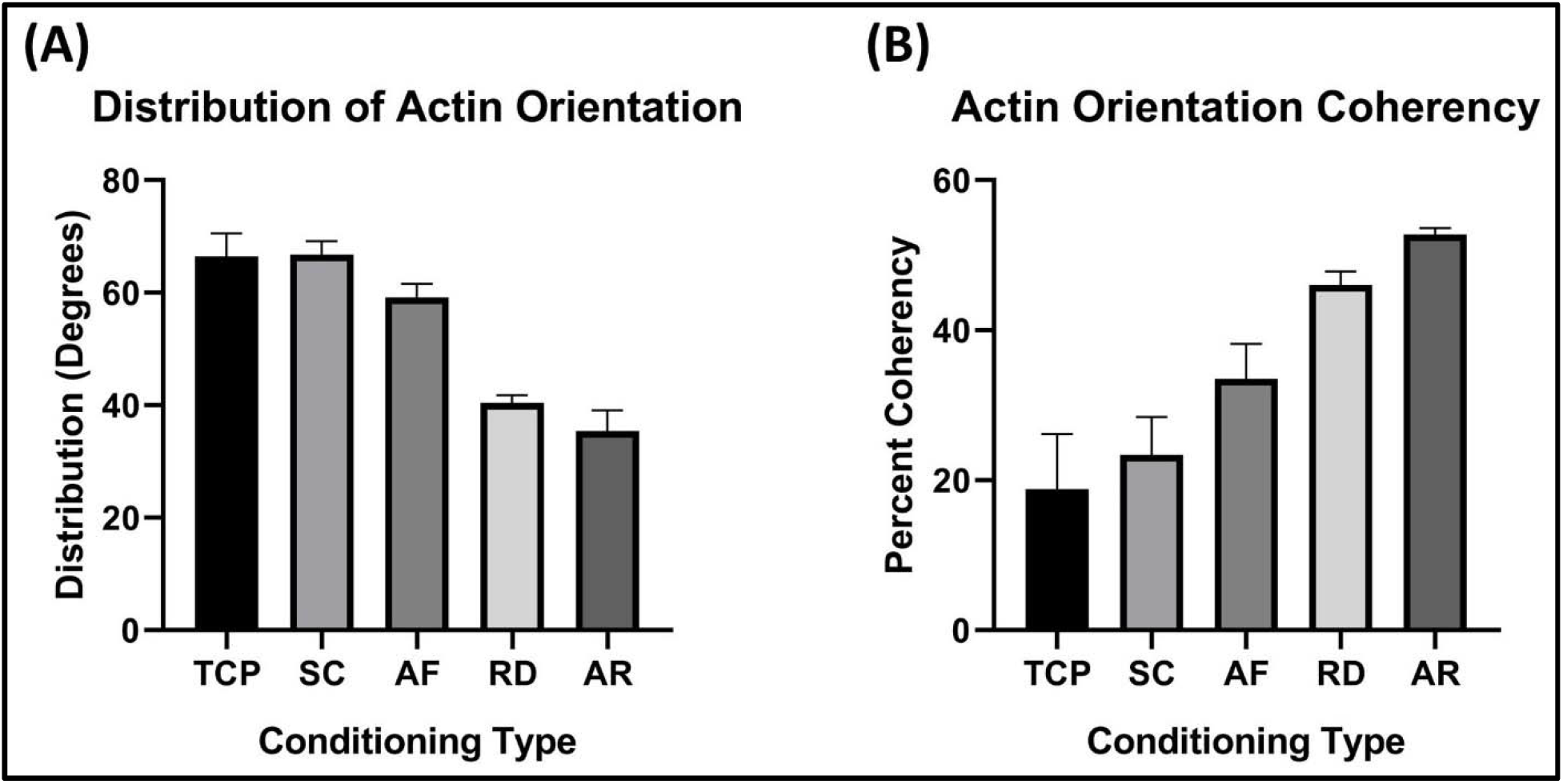
A combination of angular radial (AR) conditioning reduces actin distribution and increases orientation coherency. (A) TCP and SC control groups present large standard deviations in actin filament orientation in comparison to AF, RD, and AR conditioned groups. The actin filament of RD and AF groups express the lowest deviation, respectively. (B) Similarly, analysis of actin filament dominant orientation coherency from each group shows that TCP and SC control groups were highly disordered in comparison to the conditioned groups. Furthermore, actin filament in the AR groups were the most coherent, with the largest percentage of actin filament being aligned in the dominant orientation, n = 3.

